# A Live Attenuated Vaccine Model Confers Cross-Protective Immunity Against Different Species of *Leptospira* spp.

**DOI:** 10.1101/2020.04.13.039438

**Authors:** Elsio A. Wunder, Haritha Adhikarla, Camila Hamond, Katharine A. Owers, Li Liang, Camila B. Rodrigues, Vimla Bisht, Jarlath E. Nally, David P. Alt, Mitermayer G. Reis, Peter J. Diggle, Philip L. Felgner, Albert I. Ko

## Abstract

Leptospirosis is the leading zoonotic disease in terms of morbidity and mortality worldwide with a great health and economic impact in both humans and animals. Effective prevention is urgently needed as rapid urbanization, climate change and drivers of disease transmission continue to intensify. The key challenge has been developing a widely-applicable vaccine that protects against the 13 different pathogenic species and >300 serovars that can cause leptospirosis, providing a major public health benefit and opportunity to leverage One Health approaches. Live attenuated mutants that can boost immunity and induce protection are enticing vaccine candidates and poorly explored in the field. We evaluated a recently characterized motility-deficient mutant lacking the expression of a flagellar protein, FcpA. Although the *fcpA*^-^ mutant has lost its ability to penetrate mucous membranes and cause disease, a transient bacteremia prior to clearance by the host immune response was observed. In two animal models, immunization with a single dose of the *fcpA*^-^ mutant was sufficient to induce robust anti-protein antibodies response that promoted protection against infection with different *Leptospira* spp. species. Furthermore, characterization of the immune response identified a small repertoire of biologically relevant proteins that are highly conserved among pathogenic *Leptospira* species and potential correlates of cross-protective immunity.

## INTRODUCTION

Leptospirosis is caused by a genetically and antigenically diverse group of spirochetes of the *Leptospira* genus ^1-3^. Currently the *Leptospir*a genus comprises 64 species with more than 300 serovars, with 17 of those species containing strains that can potentially cause severe disease in humans and animals ^4,5^. A broad range of mammalian reservoirs harbor the spirochete in their renal tubules shedding the bacteria in their urine for long periods of time ^6,7^. Leptospirosis is an environmentally-transmitted disease, for which the primary mode of transmission to humans is through contact with contaminated water or soil ^8^.

Although *Leptospira* has a worldwide distribution, the large majority of the burden occurs in the world’s most impoverished populations ^9^, where the rapid growth of urban slums worldwide has created conditions for rat-borne transmission. The disease causes life-threatening manifestations such as Weil’s disease ^3,10^ and pulmonary hemorrhage syndrome (LPHS) ^11^. A recent study estimated that leptospirosis causes 1.03 million cases and 58,900 deaths each year ^9^. Case fatality for Weil’s disease and LPHS is >10 and >50%, respectively, despite aggressive supportive care ^11^. These estimates place leptospirosis as a leading zoonotic cause of morbidity and mortality worldwide. The burden of leptospirosis will increase as climate and land use change continues to evolve and the world’s slum population doubles to two billion by 2025 ^12^.

The public health priority is therefore prevention of leptospirosis before severe complications develop. However, there is no effective control for leptospirosis and safe and efficacious vaccines for are not available for human use ^1,2^. China and Cuba use whole-cell vaccines in humans ^13,14^, but they are not licensed to be used elsewhere.

Whole-cell vaccines are widely used for veterinary purposes but have significant limitations, since immunity is of short duration and predominantly humoral against LPS, which are serovar-specific moieties. Multivalent vaccines are unable to achieve sufficient coverage against the spectrum of serovars that are important for animal and human health ^3^. Research has thus focused on characterizing surface-associated and host-expressed proteins as sub-unit vaccine candidates ^2,3^. To date, these conventional approaches have not yielded candidates and attempts have failed to identify a universal, widely-applicable vaccine.

Current control measures have been uniformly ineffective in addressing the large human and animal global health burden due to leptospirosis, especially in developing countries. Given the limitations of the whole-cell vaccines available and the ineffective attempts to identify protein vaccine candidates ^3^, an attenuated vaccine approach remains a feasible strategy. Attenuation of *Leptospira* virulence has been long-recognized yet poorly-understood phenomenon ^3,15^. Until recently, the inability to produce well-defined mutants has preempted efforts to identify a safe and efficacious attenuated vaccine. However, current advances in genomic tools and whole-genome sequencing data for *Leptospira* ^1,6^ have circumvented this limitation and some promising results have been shown ^15,16^.

Recently, our group identified and characterized a novel flagellar protein in the Leptospira genus involved in the composition of the sheath of the leptospiral flagella, Flagella-coil protein A (FcpA). A mutant deficient in the *fcpA* gene lost its ability to produce translational motility and to penetrate mucous membranes, resulting in loss of kidney colonization and lethality in the hamster model of leptospirosis. Although highly attenuated in the hamster model, a needle inoculation of the mutant produced a transient bacteremia prior to clearance by the host immune response ^17^. In the present study, we evaluated the *fcpA*^-^ motility-deficient mutant as a potential candidate for a live attenuated vaccine.

## RESULTS

### A motility-deficient strain as an attenuated-vaccine candidate

We characterized a previously-unidentified flagellar sheath protein (FcpA) that was essential for translational motility and thus for virulence ^17^. Despite the phenotype of complete attenuation, we observed that the L1-130 *fcpA*^-^ mutant caused a transient systemic infection, which was cleared 7 days after intraperitoneal inoculation of 10^8^ leptospires in hamsters ^17^. In this study, after inoculation of 10^7^ leptospires using the subcutaneous route of infection in hamsters, we detected presence of DNA of the mutant by qPCR in all the tissues tested, with the exception of the brain (Figure 1A). These results were similar to those observed previously, with the wild-type reaching higher number of leptospires in all tissues analyzed, leading to the death or euthanasia of the animals due to clinical signs of disease 5-7 days after infection. In comparison, the signal for the *fcpA*^-^ mutant strain was undetectable after 7 days with all inoculated animals surviving with no detectable leptospires in either kidney or blood, measured by qPCR and culture. Similarly, no detectable signal was observed for the animals immunized with the L1-130 heat-killed strain (Figure 1A). We also tested the *fcpA*^-^ mutant in the mouse model using different doses of infection (Figure 1B). Although the dose of the wild-type strain was not enough to produce disease and lethality on infected mice, all animals were colonized and the presence of the leptospiral DNA in blood was detectable until the 15^th^ day after infection (Figure 1B). Furthermore, no dose of the *fcpA*^-^ mutant caused colonization (data not shown) and there was a significant difference in the magnitude of dissemination of the mutant in the blood compared to the wild-type (Figure 1B). DNA signal of the *fcpA*^-^ mutant was only observed in the blood of animals infected with doses of 10^7^ and 10^5^ until the 13^th^ and 8^th^ day after infection, respectively. Taken together, these results indicate that although the *fcpA*^-^ mutant is attenuated in both hamster and mouse model, there is a hematogenous dissemination of this mutant, identified by detection of its DNA. The mutant appears to be cleared by the immune system before it results in observable disease or death of the animals. Furthermore, we observed that the dissemination of the mutant is dose dependent. However, it is important to notice that although we don’t see any signal of the mutant in doses equal or lower to 10^3^ leptospires, the theoretic limit of detection of the qPCR assay used here ^17,18^, is 100 leptospires/mL of blood which can result in false negative results.

**Figure 1.**
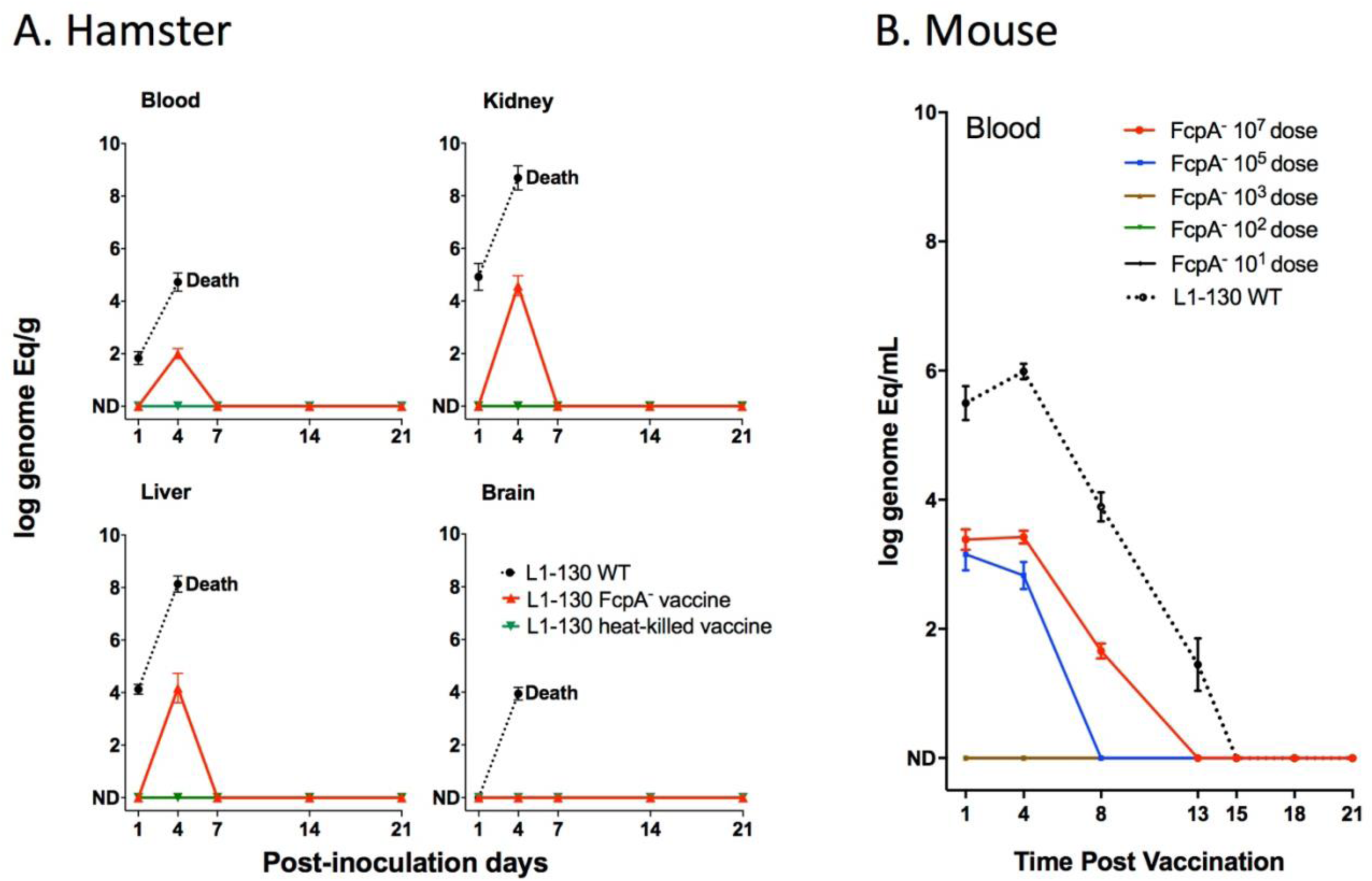
Dissemination of L1-130 FcpA^-^ mutant in animal tissues. (A) Kinetics of infection of L1-130 WT, L1-130 FcpA^-^ vaccine and L1-130 heat-killed vaccine in blood, kidney, liver and brain of hamsters after inoculation with 10^7^ bacteria. All animals infected with WT strain died between 5-6 days post-infection; (B) Kinetics of infection of L1-130 WT (10^7^ leptospires) and L1-130 FcpA^-^ attenuated vaccine (dose range from 10^7^ to 10^1^ leptospires) in blood of mouse. Results are expressed by logarithmic genome equivalent per gram or milliliter of tissues with mean and standard deviation. All doses were inoculated by subcutaneous route in both models.

### Model for cross-protective immunity to leptospirosis

We hypothesized that the transient infection produced by the *fcpA*^-^ mutant induces cross-protective responses, given previous findings ^15,17^. Immunization with the *fcpA*^-^ mutant (Figure 2A) conferred complete protection against mortality in hamsters from infection with homologous and heterologous serovars (Figure 2B and Table S2). In contrast, immunization with heat-killed leptospires conferred partial protection to the homologous but not against heterologous serovars (Figure 2B and Table S2). It’s important to mention that the strain Hardjo 203 was described to cause only colonization in the hamster model infected by intraperitoneal route ^19^. However, in our experiments using conjunctival route we reproducibly observed 25% death rate in the non-vaccinated group but no deaths in the vaccinated group, which explains the wide 95% CI range (Figure 2B and Tables S2).

**Figure 2.**
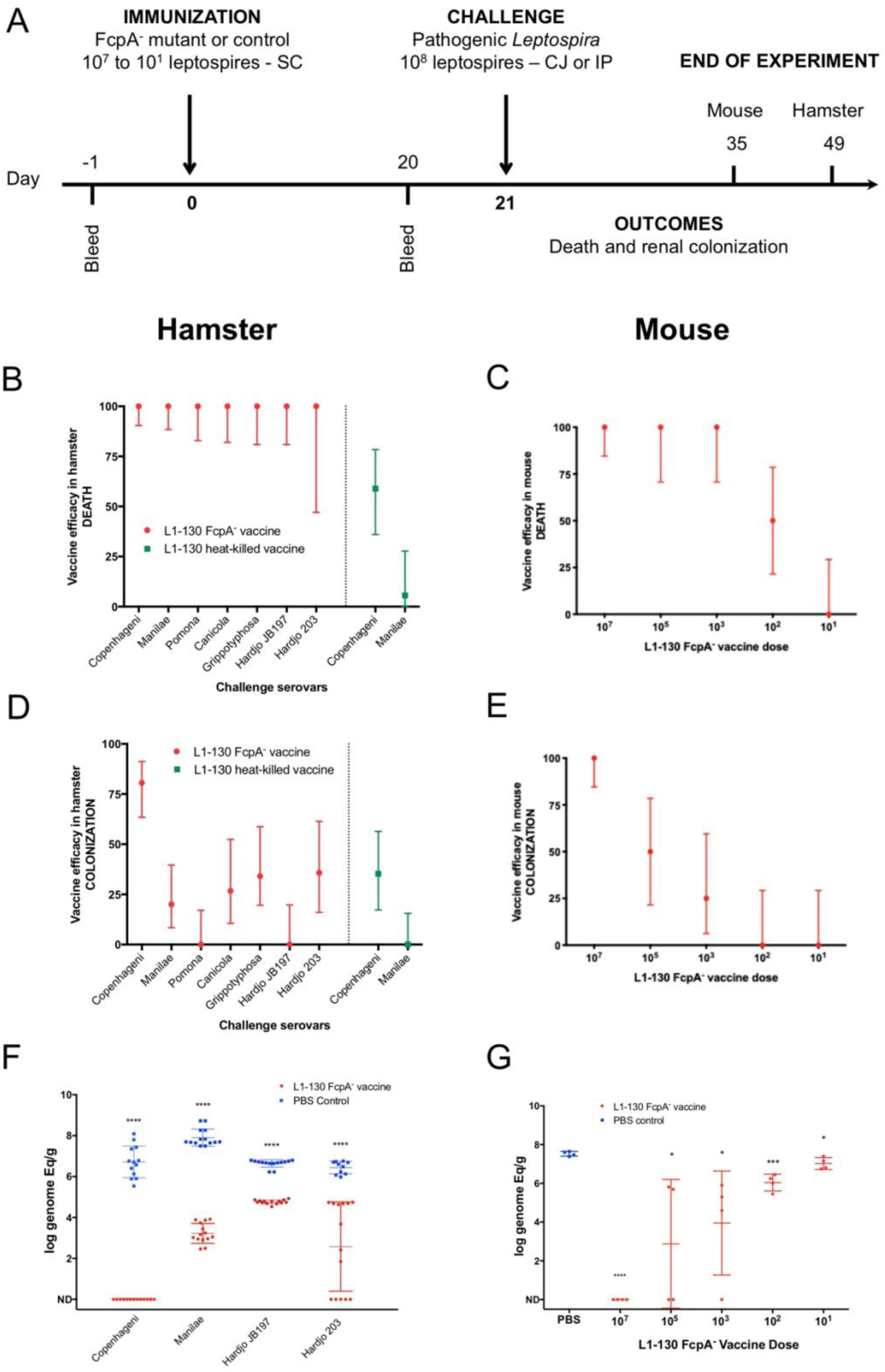
Efficacy of L1-130 FcpA^-^ attenuated-vaccine model. Animals were vaccinated with a dose of 10^7^ leptospires (hamsters) or a range of doses from 10^7^ to 10^1^ leptospires (mice) by subcutaneous route (SC). Animals were bled the day before immunization (day -1) and day 20 post-immunization (A). Hamsters were challenged with either the homologous strain or different heterologous strains. Mice were challenged with the heterologous serovar Manilae of *L. interrogans*. Efficacy of the vaccine against death and colonization was evaluated for hamsters (B and D) and mice (C and E) and represented by percentage and 95% CI based on frequency of outcomes compared to PBS-immunized animals. Hamster experiment are showing the results after vaccination with the FcpA^-^ attenuated-vaccine (red) and heat-killed vaccine (blue). Bacterial load in the kidney was measured by qPCR in hamsters (F) and mice (G) and compared between PBS-immunized animals (blue) and animals immunized with FcpA-attenuated-mutant (red). Results are expressed in logarithmic genome equivalents per gram of renal tissue with mean and standard deviation. Asterisk symbols represent statistical significance calculated by t-test: * p<0.01, *** p<0,0001. See also Supplementary tables 2 and 3.

Protection against renal colonization was only observed in 80% of the animals immunized with *fcpA*^-^ mutant after homologous infection. Heterologous infection gave varying levels of protection, from 0% to 35.7% (Figure 2D and Table S2). Hamsters are highly susceptible to leptospirosis ^20^, so the finding that the attenuated strain conferred partial protection against colonization was not unexpected. To understand the efficacy of the *fcpA*^-^ mutant vaccine to protect against colonization, we tested different doses of immunization using the mouse model against heterologous infection. Our results indicate that the protection conferred by the *fcpA*^-^ mutant is dose dependent. Against death, the vaccine conferred 100% protection up to a dose of 10^3^ leptospires of the *fcpA*^-^ mutant (Figure 2C and Table S3), but a dose as high as 10^7^ leptospires was necessary to obtain 100% protection against colonization (Figure 2E and Table S3). Furthermore, our quantitative analyzes of renal colonization showed that although the *fcpA*^-^ mutant can’t promote complete protection, there is a significant reduction of the burden of the disease both in hamster after heterologous infection (Figure 2F and Table S2) and in lower doses of the vaccine in the mouse model, which also revealed a dose dependent phenotype (Figure 2G and Table S3).

These findings indicate that a single dose of a live attenuated vaccine elicits cross-protective immunity against serovars belonging to *L. interrogans, L. kirschneri* and *L. borgpetersenii*, the species which encompasses the majority of serovars of human and animal health importance.

### Antibodies against *Leptospira* proteins as a correlate for the cross-protective immunity

The *fcpA*^-^ attenuated vaccine induced a weak agglutinating antibodies response to the homologous serovar, Copenhageni, and undetectable MAT titers against heterologous serovars, both in hamsters (Figure 3A) and mouse (Figure 3C). Furthermore, in the mouse model, agglutinating antibodies were only measurable with a dose of at least 10^5^ leptospires (Figure 3C). In contrast, a single dose of the *fcpA*^-^ mutant was able to induce a robust immune response against leptospiral proteins, recognizing proteins across the different species of *Leptospira* used in the hamster model (Figure 3B) and the heterologous strain used in the mouse model with a dose of at least 10^3^ leptospires (Figure 3D). In addition, the presence of detectable antibodies measured by ELISA correlates with the highest dose that induced 100% protection against death in the mouse model (10^3^ leptospires), and there is a decreased on the OD for all doses when the Manilae antigen was treated with proteinase K (Figure 3E). Furthermore, our passive transfer experiments using hamster-immune sera against *fcpA*^-^ attenuated-vaccine conferred 100% protection against heterologous lethal infection in hamsters (Figure 3F) and mouse (Figure 3G). Taken together, these results indicate that anti-*Leptospira* protein antibodies, and not agglutinating antibodies, are the correlate of vaccine-mediated cross-protective immunity.

**Figure 3.**
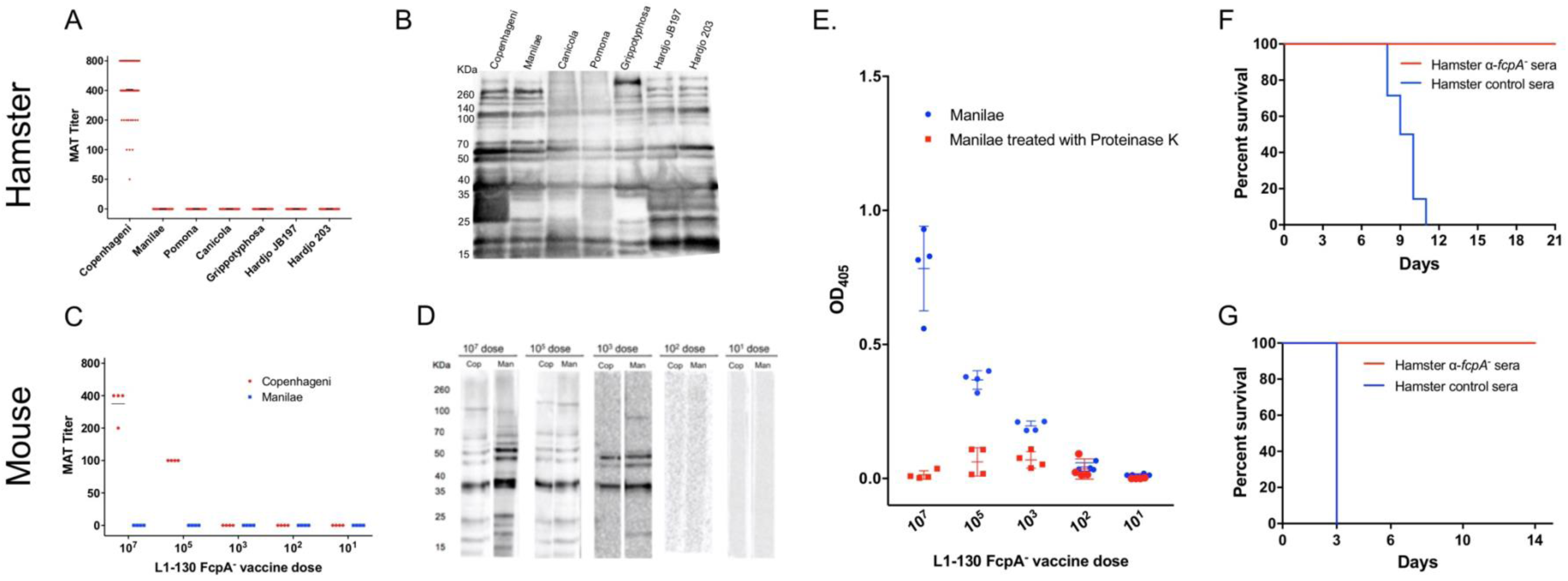
Immunogenicity and correlates of immunity for L1-130 FcpA^-^ attenuated-vaccine model. Individual sera of hamsters and mice were obtained after 20 days post-vaccination by a subcutaneous (SC) dose of 10^7^ leptospires (hamsters) or a range of doses from 10^7^ to 10^1^ leptospires (mice) of the attenuated-vaccine. MAT (A and C) and western-blot (B and D) were performed adopting as antigen all the strains used for challenged in both hamster and mice, respectively. Mice sera was additionally tested using an ELISA assay (E) adopting whole-cell extract of serovar Manilae with (red) and without (blue) Proteinase K treatment as antigen. Furthermore, a pool of hamster immune-sera vaccinated with a dose of 10^7^ leptospires of FcpA^-^ attenuated-vaccine was used for passive transfer experiments. 2-mL of sera was passively transfer to naïve hamsters (F) or mice (G) followed by challenge with a dose of 10^8^ leptospires of heterologous serovar Manilae by conjunctival (CJ) or intraperitoneal (IP) route, respectively. Results are expressed in a survival curve of animals passively transfer with FcpA^-^ anti-sera (red) and control hamster sera (blue).

### Highly-conserved seroreactive proteins as potential targets for eliciting cross-protective responses

We characterized the antibody response to the attenuated vaccine using a downsized proteome array of 660 and 330 ORFs for hamster and mouse sera, respectively. We identified a total of 133 (Figure 4A) and 56 (Figure 4B) protein targets on our analysis of hamster (Hamster 10^7^) and mouse (Mouse 10^7^) respectively, immunized with a dose of 10^7^ leptospires and a total of 13 protein targets (Figure 4C) on our analysis of mouse immunized with different doses of the attenuated vaccine (Mouse all). The reason to analyze the mouse results separately was based on the fact that a dose of 10^7^ leptospires of the attenuated-vaccine was able to give 100% cross-protection against lethality and colonization (Figure 2C and E). When combined, these three different analyses resulted in a total of 154 unique protein targets (Figure 4D and Table S4). Of those, 55% (85) have no prediction of localization and 23% (36), 14% (21) and 8% (12) have prediction to be Cytoplasmic membrane-associated, Outer Membrane proteins (OMP), and Cytoplasmic, respectively (Figure S1A). Enrichment analysis showed a 5.0-fold (p=4.51E-10) and 1.8-fold (p=2.92E-04) enrichment for OMP and Cytoplasmic membrane-associated, respectively(Figure S1B). In contrast, Cytoplasmic proteins were 0.3-fold (p=2.91E-10) underrepresented in reactive antigens groups (Figure S1B).

**Figure 4.**
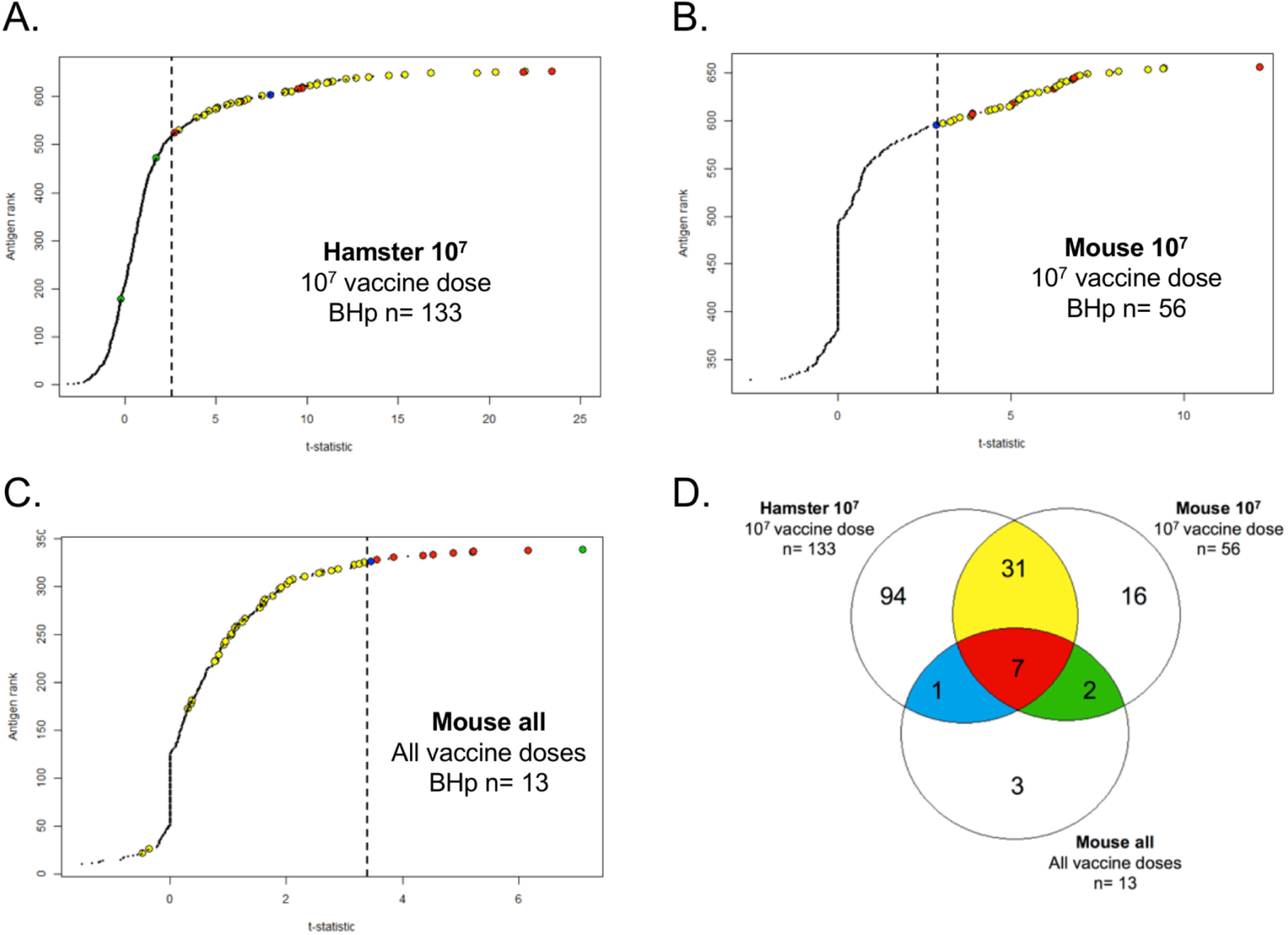
Proteome array analysis of immune-sera against L1-130 FcpA^-^ attenuated-vaccine. Using statistical modeling we calculated the t-statistics value for each individual antigen used in the proteome array (660 for hamster and 330 for mice) based on three groups: the contrast between vaccinated and unvaccinated hamsters (A) or mice (B) using a vaccine dose of 10^7^ leptospires; the dose response relationship for each antigen on mice (C) vaccinated with a range of doses from 10^7^ to 10^1^ leptospires of the attenuated vaccine. Results are ranked based on individual t-statistics values for each antigen, and the dashed line represents the selection point for the antigens based on Bhp-test. The Venn-diagram (D) shows the relationship of all the 154 antigens identified in the three groups. The subgroups of antigens selected for further characterization are highlighted in color. See Figure 5.

Clusters of Orthologous Groups of proteins (COGs) were widely represent in those targets (Table S4), with at least 1 protein in each of the 18 functional categories. The COGs with higher representation were General function prediction only (R), Cell wall/membrane/envelope biogenesis (M) and Intracellular trafficking, secretion, and vesicular transport (U) and Cell Motility (N) with 19, 17, 16 and 14 proteins, respectively. However, in the addition to the 11 protein targets assigned as Function unknown (S), the vast majority of the proteins had no COG assigned (59) (Figure S1C). Enrichment analysis identified proteins with predicted COG-U, COG-N and COG-M function as highly enriched among the reactive antigens, by 4.9-fold (p=2.27E-07), 3.1-fold (*p* = 8.35E-05), and 1.6-fold (p<0.05), respectively (Figure S1D). Furthermore, proteins predicted to be involved in signal transduction mechanisms (COG-T) and in amino acid transport and metabolism (COG-E) were significantly underrepresented in reactive antigens, by 0.4-fold (p=0.016) and 0.3-fold (0.02), respectively (Figure S1D). Taken together, the enrichment analysis validates our approach to identify biologically relevant proteins candidates for a cross-protective vaccine.

We were able to narrow down the identified 154 proteins to 41 protein targets based on their relationship among the three different groups of the proteome array’s analysis (Figure 5 and Figure S2). Seven proteins were identified in all groups (Figure 4D, Figure 5 and Table S4, red) and 31 proteins were identified in both hamster and mouse vaccinated with a dose of 10^7^ leptospires of the attenuated-vaccine (Figure 4D, Figure 5 and Table S4, yellow). Furthermore, we identified 3 extra proteins identified in the group of mice immunized with different doses, 2 between the group of mice immunized with a dose of 10^7^ leptospires (Figure 4D, Figure 5 and Table S4, green) and 1 extra protein between the group of hamsters immunized with a dose of 10^7^ leptospires (Figure 4D, Figure 5 and Table S4, blue). Hamster and mice immune sera were highly reactive to the majority of the 41 proteins (Figure 5), in contrast with the low reactivity for the control sera and animals vaccinated with the heat-killed vaccine (Figure S2), indicating the ability of the attenuated vaccine to induce immunity against leptospiral proteins. We identified plausible vaccine candidates among these 41 seroreactive proteins (Figure 5), which included 6 outer membrane proteins and known putative virulence factors such as LipL32, LipL41 and Lig proteins ^1,2^, providing supportive evidence for using proteome arrays to identify such proteins. Not surprisingly, 40% of those targets are identified as hypothetical proteins with no described function. However, the majority (70%) have high amino acid sequence identity (>80%) among their respective orthologs in all the 13 pathogenic *Leptospira* species (Figure 5), and therefore may be targets for eliciting cross-protective responses. Moreover, sera from confirmed patients with acute leptospirosis reacted with 17 of the 41 *Leptospira* proteins recognized by sera from animals immunized with the attenuated vaccine (Figure S2).

**Figure 5.**
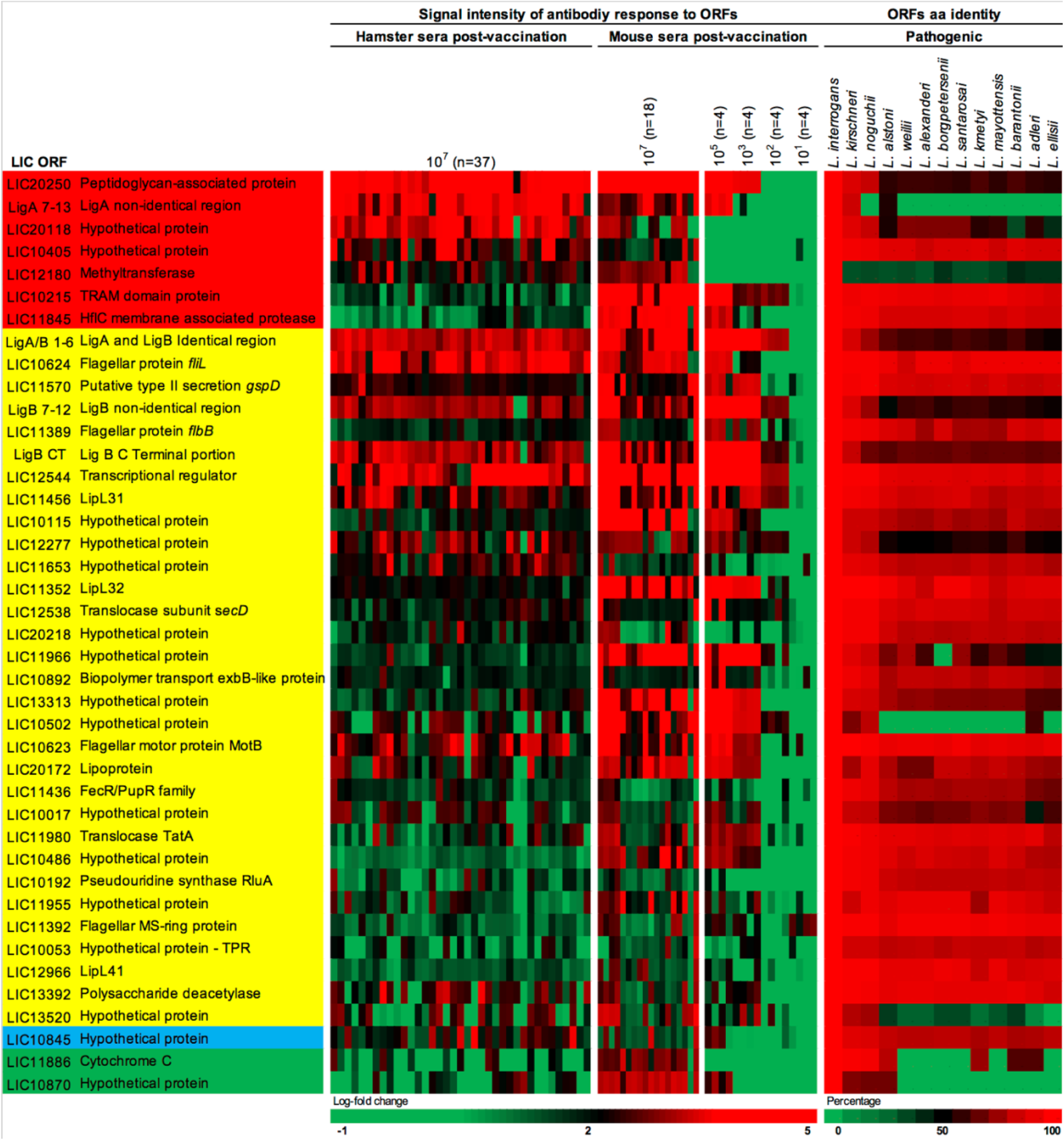
Heat-map of 41 seroreactive proteins recognized by hamsters and mice immunized with attenuated L1-130 FcpA^-^ attenuated-vaccine. Proteins were selected based on the groups depicted on Figure 4 and Table S4: present in all three groups of analysis (red), present in both hamster and mice immunized with 10^7^ leptospires (yellow), present in both hamsters immunized with 10^7^ leptospires and mice immunized with a dose range (blue), and present in both mice immunized with 10^7^ leptospires and mice immunized with a dose range (green). The proteins are identified by their *L. interrogans* serovar Copenhageni ORF number and the heat-map shows the signal intensity of antibody response (based on log-fold change) in all animals vaccinated with the *fcpA*^-^ mutant used for this analysis (37 hamsters and 34 mice). Right panel shows amino acid sequence identity of respective ORFs among a representative of all pathogenic *Leptospira* species. See Figure S2.

## DISCUSSION

Live attenuated vaccines are one of the most universal vaccination technologies used for prevention of important bacterial diseases like tuberculosis, cholera and salmonellosis and viruses like Yellow Fever, Influenza and Zoster ^21,22^. Despite the risks and the hurdles to identify the best candidates, including the best balance between attenuation and immunogenicity, research in this field continues to increase ^22,23^. An effective vaccine against leptospirosis would provide interdependent health and societal benefits by preventing transmission and disease in livestock and domestic animals and reducing the risk of spill-over infections in humans. In this proof-of concept study, we showed the efficacy of the *fcpA*^-^ attenuate-vaccine in preventing both death and renal colonization in animal models.

The transient bacteremia produced by needle inoculation of a single dose of the *fcpA*^-^ mutant was sufficient to induce robust and cross-protective immunity in two different animal models of leptospirosis. The first live-attenuated vaccine for leptospirosis with a defined mutation has been recently described ^15,16^ showing cross-protective immunity in hamsters and the potential for this approach. These authors made the important finding that immunization with an attenuated, LPS-deficient strain of serovar Manilae conferred varied levels of cross-protection against different species after one or two doses of the attenuated-vaccine. In our experiments, immunization with a single dose of the live *fcpA-*mutants conferred complete protection against death after heterologous infection on both hamster and mouse models. Furthermore, we observed that the cross-protection against death was dose dependent. In another spirochete, a similar approach has been successfully explored when a single dose of a flagella-less mutant in *Borrelia burgdorferi* induced homologous protection in mouse ^24^. More recently, a target mutant in *B. burgdorferi* induced cross immunity, but four immunizations were necessary for full protection ^25^. Previous work with attenuated-vaccine in *Leptospira* showed evidence of protection against homologous challenge in guinea-pigs, cattle, hamsters and swine, but only partial cross-protection in hamsters and gerbils ^3,15^. To our knowledge, this is the first study showing evidence of cross-protection across species with a vaccine in leptospirosis.

A single high-dose of the *fcpA*^-^ mutant induced sterilizing immunity in the mice model of leptospirosis. Other than reduction of lethality, the ability to prevent transmission is a key parameter to determine the success and feasibility of a leptospiral vaccine. Whole-cell vaccines have shown divergent levels of protection against colonization in different species ^26-29^. In our hamster model we observed a significant reduction of the bacterial burden in the kidney of vaccinated animals, without significant levels of protection. Mice models are good alternatives for sublethal leptospirosis infection and particularly to study renal colonization, compared to the highly susceptible hamster model ^30^. Previous experiments in mice showed that if the challenge dose is not able to cause death of the animals, a portion of the sublethal dose of leptospires is able to escape the blood defenses and colonize the kidneys ^31^. Furthermore, the dose of infection necessary to cause disease in natural infection remains unclear, although recent research showed that the numbers of leptospires available in the environment are low on average ^32,33^. Current animal model experiments focus on unrealistic high challenge doses, which can devaluate vaccine efficacy estimation. Despite that, our results indicate that the *fcpA*^-^ attenuated-vaccine is able to induce a robust sterilizing immunity capable of preventing death and colonization after a high lethal dose of infection.

Anti-*Leptospira* protein antibodies, and not agglutinating antibodies, are the correlate of *fcpA*^-^ attenuated vaccine-mediated cross-protective immunity. The essential role of antibodies for protection against leptospirosis has been long described ^34,35^. However, the key knowledge gap for vaccine development is whether naturally-acquired infection elicits immunity to reinfection. There have been no attempts to address this question in either humans or animals and our understanding has thus relied solely on experimental models of acquired immunity. Passive transfer experiments have shown that whole-cell vaccine immunity is antibody-mediated ^3,35^. Serovar-specific protection correlates with agglutinating antibody titers ^34,36^, which are directed against lipopolysaccharides (LPS) ^37^. Anti-leptospiral LPS antibodies are generated by whole-cell based vaccines, and the restricted protection these vaccines confer is one of their major limitations ^3^. Recently, immunization with an attenuated, LPS-deficient strain of serovar Manilae conferred protection against infection with serovar Pomona and other *L. interrogan*s species, suggesting that anti-*Leptospira* protein responses are indeed cross-protective ^15,16^. Our results not only corroborate those previous studies, but also shows strong evidence, through passive transfer experiments, that the cross-protection mediated by anti-protein antibodies is transferable.

Comprehensive high-throughput analysis of the antibody response identified potential protein candidates with a major role on the cross-protection conferred by live-attenuated vaccine. The attenuated vaccine induced a robust anti-*Leptospira* protein antibody response, but these antibodies recognized a limited repertoire of 154 antigens. Although the numbers in our study are higher compared to the 11 proteins reported by a similar study ^15^, this difference likely relates to the significantly higher sensitivity of the proteome array compared to the 2-DIGE method used previously. Nevertheless, our approach was validated by the enrichment analysis of the reactive proteins, which showed that proteins classified as OMP and cytoplasmic membrane-associated, and proteins predicted to be involved in cell wall/membrane and intracellular trafficking/secretion were highly enriched among the selected proteins. Moreover, using two animal models and different doses of the vaccine allowed us to reduce this number to 41 selected proteins, characterizing correlates of immunity. By implementing a systematic approach to identifying cross-protective antigens, we address a critical barrier to exploiting this database of genomic information for vaccine discovery.

Given the limitations of the whole-cell vaccines currently in use, identifying the perfect protein candidate has been a major effort in the field of vaccine development for leptospirosis ^3^. Immunization with Lig proteins or LipL32 ^2,38,39^ has been shown to confer protection, albeit not cross-protective immunity, against experimental infection. However, efficacy results have been proven difficult to replicate ^2,40^. As of yet we do not know whether a single protein can be immunoprotective or if the additive value of immunity against a group of proteins is essential for cross-immunity. Lately, studies have shown that a chimeric approach has the potential for an effective universal vaccine against leptospirosis ^41,42^. Other than Ligs and LipL32, LipL41 has been tested as a vaccine candidate without success ^2^ and LIC20172 and LIC11966 have been identified and tested, respectively, as promising candidates for subunit vaccine ^43,44^. More recently, a DNA vaccine delivering two leptospiral proteins (RecA and FliD) has shown promising results including high survival rates, sterilizing immunity and heterologous protection, although tested only among two different *L. interrogans* strains ^45^. A major limitation for vaccine development for leptospirosis is the lack of evidence whether immunological correlates from experimental animals are relevant to human disease. However, our results suggest that the same antigens which confer cross-protective immunity in the attenuated vaccine model may play a role in naturally-acquired immunity in humans.

This study is the first description of vaccine that elicits cross-protection against disease caused by different species of the genus *Leptospira*, and that the protection is transferable through anti-protein antibodies. Effective prevention is urgently needed as drivers of disease transmission continue to intensify, like rapid urbanization and climate change. The *fcpA*^-^ mutant induced robust and cross-protective immune responses against protein moieties conserved across pathogenic *Leptospira*. We identified a small repertoire of biologically relevant proteins that can be used in the field of vaccine and diagnostic development for leptospirosis. Characterization of these proteins as novel putative virulence factors can also provide new insights on the pathogenesis of the disease. Both approaches are currently being tested by our group as we try to identify an absolute correlate of immunity for leptospirosis. Nonetheless, these findings indicate that a single dose of *fcpA*^-^ mutant elicits cross-protective immunity against serovars belonging to species of *Leptospira* that encompasses the majority of serovars of human and animal health importance. Taken together, those findings highlight the feasibility to use this model to formulate novel approaches for prevention, such as a synergistic development of vaccines for human and animal health in a diverse range of epidemiological settings.

## ACKNOWLEDGEMENTS

We would like to thank the Leptospirosis team at Instituto Gonçalo Moniz, Oswaldo Cruz Foundation, Salvador, Brazil, for all the support and technical assistance. This work was supported by NIH grants R01AI052473, U01AI088752, R25TW009338, R01TW009504, and R01AI121207 (A.I.K,). C.H. was supported by Programa Ciências sem fronteiras, CNPq, Brazil.

## AUTHOR CONTRIBUTIONS

E.A.W.J., H.A. and A.I.K. designed the experiments, E.A.W.J., H.A., C.H., L.L., C.B.R., and V.B. performed the experiments. E.A.W.J., H.A., K.O., L.L., P.J.D., and P.L.F. performed the data analysis. M.G.R., J.E.N. and D.P.A. contributed key reagents. E.A.W.J. and H.A. wrote the original draft of the manuscript with the other authors providing comments and edits to the final version.

## METHODS

### Vaccine and challenge strains

Leptospires were cultivated in liquid EMJH medium ^46^ supplemented with 1% rabbit serum. *Leptospira interrogans* serovar Copenhageni strain Fiocruz L1-130 *fcpA*^-^ mutant ^17^ and all the 7 different strains used for the challenge experiments (Table S1) were incubated up to 7 days at 29°C, till they reached log phase (between 4-5 days of culture). For all immunization or infection experiments, the correct number of *Leptospira* was determined by a Petroff-Hausser counting chamber (Fisher Scientific).

The heat-killed vaccine was prepared by heat-inactivating preparations of *L. interrogans* strain Fiocruz L1-130 at 56°C for 20 minutes.

### Animal experimentation

#### Dissemination studies

For the dissemination experiments with the Fiocruz L1-130 *fcpA*^-^ mutant and L1-130 heat-killed vaccine in hamsters, a group of fifteen 3-week-old male Golden Syrian hamsters (Envigo) were inoculated subcutaneously with a dose of 10^7^ leptospires in 0.5 mL of EMJH medium. A group of 3 animals were euthanized at 1, 4, 7, 14 and 21 days after infection. As a control, a group of 9 animals were infected with Fiocruz L1-130 WT using the same route and dose, and animals were euthanized at days 1 and 4 after infection. The final group was euthanized at onset of disease. After euthanizing the animals, blood, kidney, liver and brain were carefully removed, collected into cryotubes and immediately placed into liquid nitrogen before being stored at -80°C until extraction of DNA. Kidney and blood were inoculated in EMJH for culture of leptospires when necessary.

For the experiment in mice, groups of three 4-week-old female C57BL/6 mice (Jackson laboratory) were inoculated subcutaneously with different doses of the vaccine (10^7^, 10^5^, 10^3^, 10^2^, 10^1^) and a control groups with 3 animals was inoculated with Fiocruz L1-130 WT with a dose of 10^7^ leptospires. Blood was collected by retro-orbital bleeding at 1, 4, 8, 13, 15, 18 and 21 days after infection.

#### Immunization and challenge

All vaccination experiments (Figure 2A) were performed using 3-week-old male Golden Syrian hamsters (Envigo) or 4-week-old female C57BL/6 mice (Jackson laboratory), divided in groups of 6-9 or 4-8 animals, respectively. Animals were vaccinated with Fiocruz L1-130 *fcpA*^-^ mutant using the subcutaneous route. Hamsters were vaccinated with a single dose of 10^7^ leptospires and mice were vaccinated with a range of doses (10^7^, 10^5^, 10^3^, 10^2^, 10^1^) in 500 and 200 μL of EMJH medium, respectively. The heat-killed vaccine was used in a single dose of 10^7^ leptospires by subcutaneous route as a control group in hamster. In addition, groups of animals were injected with PBS (Phosphate buffered saline) and served as unvaccinated controls. Blood samples were collected the day before and 20 days post-immunization by retro-orbital bleeding.

Animals were challenged on day 21 post-immunization. Hamsters were challenged by conjunctival inoculation, which mimics the natural route of infection ^47,48^ using a lethal dose (10^8^ leptospires) of a range of serovars whose virulence has been well-characterized in our laboratory (Table S1). Hamsters vaccinated with the heat-killed vaccine were only challenged with Fiocruz L1-130 (homologous) or Manilae L495 (heterologous) strains. Mice were challenged intraperitoneally with *L. interrogans* serovar Manilae L495 (10^8^ leptospires) ^30,31^. After euthanizing the animals, kidneys were collected and stored as described above.

#### Passive transfer experiments

Immune sera against Fiocruz L1-130 *fcpA*^-^ mutant was generated by immunizing a group of ten 3-weeks old male Golden Syrian hamsters using the same protocol, described above. A group of 10 animals injected with PBS were used to obtained control sera. Animals were euthanized at day 21 post-immunization by inhalation of CO^2^. Blood was obtained by cardiac puncture, followed by separation of sera that was subsequently pooled as immune (hamster α-fcpA^-^) and control sera.

Immune or control sera were passively transfer to groups of 5 naïve female mice and 7 naïve male hamsters (6-7-week-old) in a dose of 500 and 2,000 μL, respectively, using the intraperitoneal route. After 24 hours mice and hamsters were challenged with 10^8^ leptospires *of* serovar Manilae L495 (heterologous strain) by intraperitoneal and conjunctival route, respectively, as described above.

#### Ethical statement

All animal protocols were approved by the Institutional Committee for the Use of Experimental Animals, Yale University (protocol # 2017-11424). Hamsters and mice were monitored twice daily for endpoints including signs of disease and death, up to 28-days and 14-days post-infection, respectively. Surviving animals at the end of the experiment or moribund animals presenting with difficulty moving, breathing or signs of bleeding or seizure were immediately sacrificed by inhalation of CO^2^. Before each blood collection animals were anesthetized by an open-drop method with a mixture of 20% v/v isoflurane in propylene glycol.

#### Serology

Pre- and post-vaccination sera were obtained by centrifugation of clotted blood at 1,000g for 15 min at room temperature. Sera samples were kept frozen at -20°C until analysis for the presence of antibodies against leptospires by microscopic agglutination test (MAT), ELISA, immunoblotting and proteome array.

MAT was performed using standard practices and as previously described (Ko et al, 1999, Terpstra, 2003). Serum was diluted at 1:100 and tested against all the strains used in this project (Table S1). Positives samples were subsequently titrated.

For the ELISA, whole cell lysate was prepared by centrifugation of *L. interrogans* serovar Manilae L495 and Fiocruz L1-130 cultures (10^8^ cells) at 12,000 rpm, 4°C for 20 minutes. The pellets were washed twice with Phosphate Buffered Saline (PBS) and resuspended in 500 µL of PBS. Resuspended cultures were sonicated in ice for 6 cycles at 30 kHz with a power output of 300W. Lysates were quantitated by Bradford assay and employed as antigen at a concentration of 150 ng/well (in 0.05 M carbonate buffer, pH 9.6). Flat-bottomed polystyrene microtiter plates (Corning) were coated with *Leptospira* antigen and incubated overnight at 4°C. The plates were washed three times with PBS-0.05% (vol/vol) Tween 20 (PBST) and incubated with blocking solution (5% blocking milk in 2% [wt/vol] bovine serum albumin) for 2 h at 37°C. After four washes with PBST, wells were incubated with mouse immune sera, diluted 100-fold in 2% BSA, for 1 h at 37°C. Secondary anti-mouse HRP conjugated antibody (Jackson ImmunoResearch) was used at a dilution of 50,000 (2% BSA) and incubated for 1 h at 37°C. TMB (SureBlue Reserve) was used for detection and the reaction was stopped by adding 100 μL of 2 N H^2^SO^4^. Absorbance (450 nm) was recorded by microplate reader (Biotek).

To evaluate the effect of proteolytic enzyme treatment on *Leptospira* antigen we used the protocol previously described ^49^. Briefly, *Leptospira* antigen coated in assay wells was treated with 0.1 mg of Proteinase K (Invitrogen) at 37°C for 2 h. The plates were washed three times with PBST to remove unbound proteins and followed by blocking and testing as described above.

#### qPCR

DNA was extracted from blood and tissue samples using the Maxwell®16 (Promega Corporation) instrument following the manufacturer’s instructions. Quantitative Real-time PCR assays were performed on hamster and mouse tissues using an ABI 7500 instrument (Applied Biosystems) and Platinum Quantitative PCR Supermix-UDG (Invitrogen Corporation) with *lipL32* primers and probe as described previously ^47^.

#### Western Blot

Immunoblots with whole cell extract of *Leptospira* strains were performed as previously described ^50^. Western blot was performed with a pooled of hamster or mice immune sera α-fcpA^-^ at dilution of 1:100. For subsequent detection, HRP goat anti-mouse or anti-hamster’s serum (Jackson ImmunoResearch) was employed at dilution of 1:100,000. Blots were analyzed using ChemiDoc™ Imager (Bio-Rad).

#### Proteome array

The full ORFeome was amplified from *Leptospira interrogans* serovar Copenhageni strain Fiocruz L1-130 as previously described ^51,52^. The ORFs larger than 150 bp were amplified from genomic DNA, followed by recombination cloning into a T7 expression vector. Genes larger than 3 kb were cloned as segments. A list of 660 most reactive antigens were selected from previous studies with human sera of patients with leptospirosis ^51-53^ and used for the hamster experiments. Mouse sera were tested in an array containing 330 proteins selected based on the latter. Proteins were expressed in the *in vitro* transcription/translation (IVTT) RTS 100 *E. coli* HY system (5 PRIME) and synthesized crude proteins were printed on 3-pad nitrocellulose-coated AVID slides (Grace Bio-Labs) using a Gene Machine OmniGrid 100 microarray printer (Genomic Solutions). In addition to IVTT expressed proteins, each array contained no DNA control spots consisting of IVTT reactions without the addition of a plasmid, serial dilutions of purified IgG/ spots.

The arrays were probed for IgG reactivity. For serum samples, the arrays were probed at 1/100 dilution in protein array blocking buffer (GVS) supplemented with *E. coli* lysate (Genscript) at final concentration of 10 mg/mL to block anti-*E. coli* antibodies. The arrays were incubated overnight at 4°C with constant agitation. After the incubation overnight, the arrays were washed three times with T-TBS and then incubated for 45 min at RT with biotin-conjugated anti-human IgG secondary antibody (Jackson ImmunoResearch), diluted at 1/400 in array blocking buffer, followed by Qdot® 800 streptavidin conjugate (ThermoFisher Scientific). The arrays were air dried after brief centrifugation. IgG signals were detected with ArrayCam 400-S Microarray Imaging System (Grace Bio-Labs) for Q800. The array signal intensities were quantified using QuantArray software. Mean pixel intensities are corrected for local background of all spots. Protein expression was validated by microarray using monoclonal anti-polyhistidine (clone His-1, Sigma).

In addition, 30 sera (acute and convalescent) of human patients from Salvador, Brazil with confirmed acute leptospirosis were probed in the array containing 660 proteins as described above. Patient samples were collected and select as previously described ^51^.

### Data analysis

We analyzed the log^10^ fold change (LFC) between pre- and post-vaccination proteome signal intensities. We subtracted the chip background and set the negative values to one (to avoid issues taking logarithms) before calculating the LFC. Analyses were conducted on three datasets: the hamster data, which used a single attenuated vaccine dose of 10^7^ leptospires (Hamster 10^7^), the mouse dose response data including all vaccine doses 10^1^, 10^2^, 10^3^, 10^5^, and 10^7^ (Mouse all), and the subset of mice given a dose of 10^7^ leptospires of the attenuated vaccine (Mouse 10^7^).

Exploratory analysis of mouse data showed a dose response relationship, with increased vaccine dose associated with increased mean signal intensity (Figure S3) as well as decreased death and colonization. We used a model that allowed us to quantify this dose response relationship when present and to instead measure the contrast between vaccinated and unvaccinated animals if only a single dose was used. Each antigen was modeled separately. For each antigen, we fit a linear model for the LFC in each animal *A* as *LFC*_*A*_ = *Experiment*_*A*_ + *V*_*A*_ + *LogDose*_*A*_ where experiment was a factor on four levels for the mice and two levels for the hamsters. *V*_*A*_ is an indicator variable for whether the animal *A* received the attenuated vaccine or a control injection. These terms were included in all models. The *LogDose* term was only included in the analysis of the dose response relationship in mice and is the logarithm of the dose (0 for control animals, 1, 2, 3, 5, 7). The indicator variable *V*_*A*_ prevents *LogDose* = 0 for control animals from being treated as a true zero. Our statistic of interest was the t-statistic. For the Hamster and Mouse 10^7^ models we interpreted the V_A_ t-statistics, and for the Mouse All dose response model we interpreted *LogDose* t-statistics. We used the Benjamini-Hochberg (BHp) correction ^54^ to control the false discovery rate at 0.05. This analysis was conducted in R software (R Core Team, 2013).

Prism 8 (GraphPad Software) was employed for all the statistical analysis of *in vivo* data. Fisher’s exact test and analysis of variance (ANOVA) were applied to assess statistical differences between pairs of groups and multiple groups, respectively. A *P* value of <0.05 was considered significant. Protein homologies of *L. interrogans* serovar Copenhageni strain Fiocruz L1-130 were identified by a BLAST search (http://www.ncbi.nlm.nih.gov/BLAST/). Clusters of orthologs groups (COGs), pSortB localization, transmembrane domains (TMhmm), and signal peptide (SignalP) information was obtained from Genoscope platform (http://www.genoscope.cns.fr/agc/microscope/home/). *P* value for enrichment statistical analysis was calculated using Fisher’s Exact test in the R environment (http://www.r-project.org).

## SUPPLEMENTAL INFORMATION

Supplemental information includes 3 figures and 4 tables.

## REFERENCES

1 Picardeau, M. Virulence of the zoonotic agent of leptospirosis: still terra incognita? Nature reviews. Microbiology 15, 297–307, doi: 10.1038/nrmicro.2017.5 (2017).

2 Ko, A. I., Goarant, C. & Picardeau, M. Leptospira: the dawn of the molecular genetics era for an emerging zoonotic pathogen. Nature reviews. Microbiology 7, 736–747, doi: 10.1038/nrmicro2208 (2009).

3 Adler, B. Vaccines against leptospirosis. Current topics in microbiology and immunology 387, 251–272, doi: 10.1007/978-3-662-45059-8_10 (2015).

4 Casanovas-Massana, A. et al. Leptospira yasudae sp. nov. and Leptospira stimsonii sp. nov., two new species of the pathogenic group isolated from environmental sources. International journal of systematic and evolutionary microbiology, doi: 10.1099/ijsem.0.003480 (2019).

5 Vincent, A. T. et al. Revisiting the taxonomy and evolution of pathogenicity of the genus Leptospira through the prism of genomics. PLoS neglected tropical diseases 13, e0007270, doi: 10.1371/journal.pntd.0007270 (2019).

6 Thibeaux, R. et al. Seeking the environmental source of Leptospirosis reveals durable bacterial viability in river soils. PLoS neglected tropical diseases 11, e0005414, doi: 10.1371/journal.pntd.0005414 (2017).

7 Xu, Y. et al. Whole genome sequencing revealed host adaptation-focused genomic plasticity of pathogenic Leptospira. Sci Rep 6, 20020, doi: 10.1038/srep20020 (2016).

8 Casanovas-Massana, A. et al. Quantification of Leptospira interrogans Survival in Soil and Water Microcosms. Applied and environmental microbiology 84, doi: 10.1128/AEM.00507-18 (2018).

9 Costa, F. et al. Global Morbidity and Mortality of Leptospirosis: A Systematic Review. PLoS neglected tropical diseases 9, e0003898, doi: 10.1371/journal.pntd.0003898 (2015).

10 Ko, A. I., Galvao Reis, M., Ribeiro Dourado, C. M., Johnson, W. D., Jr. & Riley, L. W. Urban epidemic of severe leptospirosis in Brazil. Salvador Leptospirosis Study Group. Lancet 354, 820–825 (1999).

11 Gouveia, E. L. et al. Leptospirosis-associated severe pulmonary hemorrhagic syndrome, Salvador, Brazil. Emerging infectious diseases 14, 505–508, doi: 10.3201/eid1403.071064 (2008).

12 UN-Habitat. The challenge of the slums: Global report on human settlements 2003. 310 (2003).

13 Yan, Y. et al. An evaluation of the serological and epidemiological effects of the outer envelope vaccine to leptospira. Journal of the Chinese Medical Association: JCMA 66, 224–230 (2003).

14 Martinez, R. et al. [Efficacy and safety of a vaccine against human leptospirosis in Cuba]. Revista panamericana de salud publica = Pan American journal of public health 15, 249–255 (2004).

15 Srikram, A. et al. Cross-protective immunity against leptospirosis elicited by a live, attenuated lipopolysaccharide mutant. The Journal of infectious diseases 203, 870–879, doi: 10.1093/infdis/jiq127 (2011).

16 Murray, G. L., Simawaranon, T., Kaewraemruaen, C., Adler, B. & Sermswan, R. W. Heterologous protection elicited by a live, attenuated, Leptospira vaccine. Veterinary microbiology 223, 47–50, doi: 10.1016/j.vetmic.2018.07.018 (2018).

17 Wunder, E. A., Jr. et al. A novel flagellar sheath protein, FcpA, determines filament coiling, translational motility and virulence for the Leptospira spirochete. Molecular microbiology 101, 457–470, doi: 10.1111/mmi.13403 (2016).

18 Stoddard, R. A., Gee, J. E., Wilkins, P. P., McCaustland, K. & Hoffmaster, A. R. Detection of pathogenic Leptospira spp. through TaqMan polymerase chain reaction targeting the LipL32 gene. Diagnostic microbiology and infectious disease 64, 247–255, doi: 10.1016/j.diagmicrobio.2009.03.014 (2009).

19 Zuerner, R. L., Alt, D. P. & Palmer, M. V. Development of chronic and acute golden Syrian hamster infection models with Leptospira borgpetersenii serovar Hardjo. Veterinary pathology 49, 403–411, doi: 10.1177/0300985811409252 (2012).

20 Haake, D. A. Hamster model of leptospirosis. Current protocols in microbiology Chapter 12, Unit 12E 12, doi: 10.1002/9780471729259.mc12e02s02 (2006).

21 Minor, P. D. Live attenuated vaccines: Historical successes and current challenges. Virology 479-480, 379–392, doi: 10.1016/j.virol.2015.03.032 (2015).

22 Detmer, A. & Glenting, J. Live bacterial vaccines-a review and identification of potential hazards. Microbial cell factories 5, 23, doi: 10.1186/1475-2859-5-23 (2006).

23 Loessner, H. et al. Improving live attenuated bacterial carriers for vaccination and therapy. Int J Med Microbiol 298, 21–26, doi: 10.1016/j.ijmm.2007.07.005 (2008).

24 Sadziene, A., Thompson, P. A. & Barbour, A. G. A flagella-less mutant of Borrelia burgdorferi as a live attenuated vaccine in the murine model of Lyme disease. The Journal of infectious diseases 173, 1184–1193 (1996).

25 Hahn, B. L., Padmore, L. J., Ristow, L. C., Curtis, M. W. & Coburn, J. Live Attenuated Borrelia burgdorferi Targeted Mutants in an Infectious Strain Background Protect Mice from Challenge Infection. Clinical and vaccine immunology: CVI 23, 725–731, doi: 10.1128/CVI.00302-16 (2016).

26 Sonada, R. B. et al. Efficacy of leptospiral commercial vaccines on the protection against an autochtonous strain recovered in Brazil. Brazilian journal of microbiology: [publication of the Brazilian Society for Microbiology] 49, 347–350, doi: 10.1016/j.bjm.2017.06.008 (2018).

27 Murray, G. L. et al. Evaluation of 238 antigens of Leptospira borgpetersenii serovar Hardjo for protection against kidney colonisation. Vaccine 31, 495–499, doi: 10.1016/j.vaccine.2012.11.028 (2013).

28 Clough, W. J., Little, P. R., Hodge, A., Chapman, V. C. & Holz, D. K. Protection of sheep by vaccination against experimental challenge with Leptospira borgpetersenii serovar Hardjo and L. interrogans serovar Pomona. New Zealand veterinary journal 66, 138–143, doi: 10.1080/00480169.2018.1441078 (2018).

29 Bolin, C. A. & Alt, D. P. Use of a monovalent leptospiral vaccine to prevent renal colonization and urinary shedding in cattle exposed to Leptospira borgpetersenii serovar hardjo. American journal of veterinary research 62, 995–1000 (2001).

30 Gomes-Solecki, M., Santecchia, I. & Werts, C. Animal Models of Leptospirosis: Of Mice and Hamsters. Frontiers in immunology 8, 58, doi: 10.3389/fimmu.2017.00058 (2017).

31 Ratet, G. et al. Live imaging of bioluminescent leptospira interrogans in mice reveals renal colonization as a stealth escape from the blood defenses and antibiotics. PLoS neglected tropical diseases 8, e3359, doi: 10.1371/journal.pntd.0003359 (2014).

32 Casanovas-Massana, A. et al. Spatial and temporal dynamics of pathogenic Leptospira in surface waters from the urban slum environment. Water Res 130, 176–184, doi: 10.1016/j.watres.2017.11.068 (2018).

33 Ganoza, C. A. et al. Determining risk for severe leptospirosis by molecular analysis of environmental surface waters for pathogenic Leptospira. PLoS medicine 3, e308, doi: 10.1371/journal.pmed.0030308 (2006).

34 Adler, B., Faine, S., Muller, H. K. & Green, D. E. Maturation of humoral immune response determines the susceptibility of guinea-pigs to leptospirosis. Pathology 12, 529–538 (1980).

35 Inada, R., Ido, Y., Hoki, R., Kaneko, R. & Ito, H. The Etiology, Mode of Infection, and Specific Therapy of Weil’s Disease (Spirochaetosis Icterohaemorrhagica). J Exp Med 23, 377–402 (1916).

36 Chapman, A. J., Everard, C. O., Faine, S. & Adler, B. Antigens recognized by the human immune response to severe leptospirosis in Barbados. Epidemiology and infection 107, 143–155 (1991).

37 Challa, S., Nally, J. E., Jones, C. & Sheoran, A. S. Passive immunization with Leptospira LPS-specific agglutinating but not non-agglutinating monoclonal antibodies protect guinea pigs from fatal pulmonary hemorrhages induced by serovar Copenhageni challenge. Vaccine 29, 4431–4434, doi: 10.1016/j.vaccine.2011.04.041 (2011).

38 Conrad, N. L. et al. LigB subunit vaccine confers sterile immunity against challenge in the hamster model of leptospirosis. PLoS neglected tropical diseases 11, e0005441, doi: 10.1371/journal.pntd.0005441 (2017).

39 Humphryes, P. C. et al. Vaccination with leptospiral outer membrane lipoprotein LipL32 reduces kidney invasion of Leptospira interrogans serovar canicola in hamsters. Clinical and vaccine immunology: CVI 21, 546–551, doi: 10.1128/CVI.00719-13 (2014).

40 Lucas, D. S. et al. Recombinant LipL32 and LigA from Leptospira are unable to stimulate protective immunity against leptospirosis in the hamster model. Vaccine 29, 3413–3418, doi: 10.1016/j.vaccine.2011.02.084 (2011).

41 Lin, X. et al. Chimeric epitope vaccine against Leptospira interrogans infection and induced specific immunity in guinea pigs. BMC microbiology 16, 241, doi: 10.1186/s12866-016-0852-y (2016).

42 Santos, J. C. & Nascimento, A. L. T. Chimeras could help in the fight against leptospirosis. Elife 7, doi: 10.7554/eLife.34087 (2018).

43 Lata, K. S. et al. Exploring Leptospiral proteomes to identify potential candidates for vaccine design against Leptospirosis using an immunoinformatics approach. Sci Rep 8, 6935, doi: 10.1038/s41598-018-25281-3 (2018).

44 Oliveira, T. L. et al. LemA and Erp Y-like recombinant proteins from Leptospira interrogans protect hamsters from challenge using AddaVax as adjuvant. Vaccine 36, 2574–2580, doi: 10.1016/j.vaccine.2018.03.078 (2018).

45 Raja, V. et al. Heterologous DNA prime-protein boost immunization with RecA and FliD offers cross-clade protection against leptospiral infection. Sci Rep 8, 6447, doi: 10.1038/s41598-018-24674-8 (2018).

46 Johnson, R. C. & Harris, V. G. Differentiation of pathogenic and saprophytic letospires. I. Growth at low temperatures. Journal of bacteriology 94, 27–31 (1967).

47 Wunder, E. A., Jr. et al. Real-Time PCR Reveals Rapid Dissemination of Leptospira interrogans after Intraperitoneal and Conjunctival Inoculation of Hamsters. Infection and immunity 84, 2105–2115, doi: 10.1128/IAI.00094-16 (2016).

48 Adhikarla, H. et al. Lvr, a Signaling System That Controls Global Gene Regulation and Virulence in Pathogenic Leptospira. Frontiers in cellular and infection microbiology 8, 45, doi: 10.3389/fcimb.2018.00045 (2018).

49 Udaykumar & Saxena, R. K. Antigenic epitopes on Mycobacterium tuberculosis recognized by antibodies in tuberculosis and mouse antisera. FEMS microbiology immunology 3, 7–12, doi: 10.1111/j.1574-6968.1991.tb04156.x (1991).

50 Lourdault, K., Cerqueira, G. M., Wunder, E. A., Jr. & Picardeau, M. Inactivation of clpB in the pathogen Leptospira interrogans reduces virulence and resistance to stress conditions. Infection and immunity 79, 3711–3717, doi: 10.1128/IAI.05168-11 (2011).

51 Lessa-Aquino, C. et al. Identification of seroreactive proteins of Leptospira interrogans serovar copenhageni using a high-density protein microarray approach. PLoS neglected tropical diseases 7, e2499, doi: 10.1371/journal.pntd.0002499 (2013).

52 Lessa-Aquino, C. et al. Proteomic features predict seroreactivity against leptospiral antigens in leptospirosis patients. Journal of proteome research 14, 549–556, doi: 10.1021/pr500718t (2015).

53 Lessa-Aquino, C. et al. Distinct antibody responses of patients with mild and severe leptospirosis determined by whole proteome microarray analysis. PLoS neglected tropical diseases 11, e0005349, doi: 10.1371/journal.pntd.0005349 (2017).

54 Benjamini, Y. & Hochberg, Y. Controlling the False Discovery Rate - a Practical and Powerful Approach to Multiple Testing. J R Stat Soc B 57, 289–300 (1995).

