## Supplementary Figures for "A Live Attenuated Vaccine Model Confers Cross-Protective Immunity Against Different Species of *Leptospira* spp."

**Supplementary Figure 1**

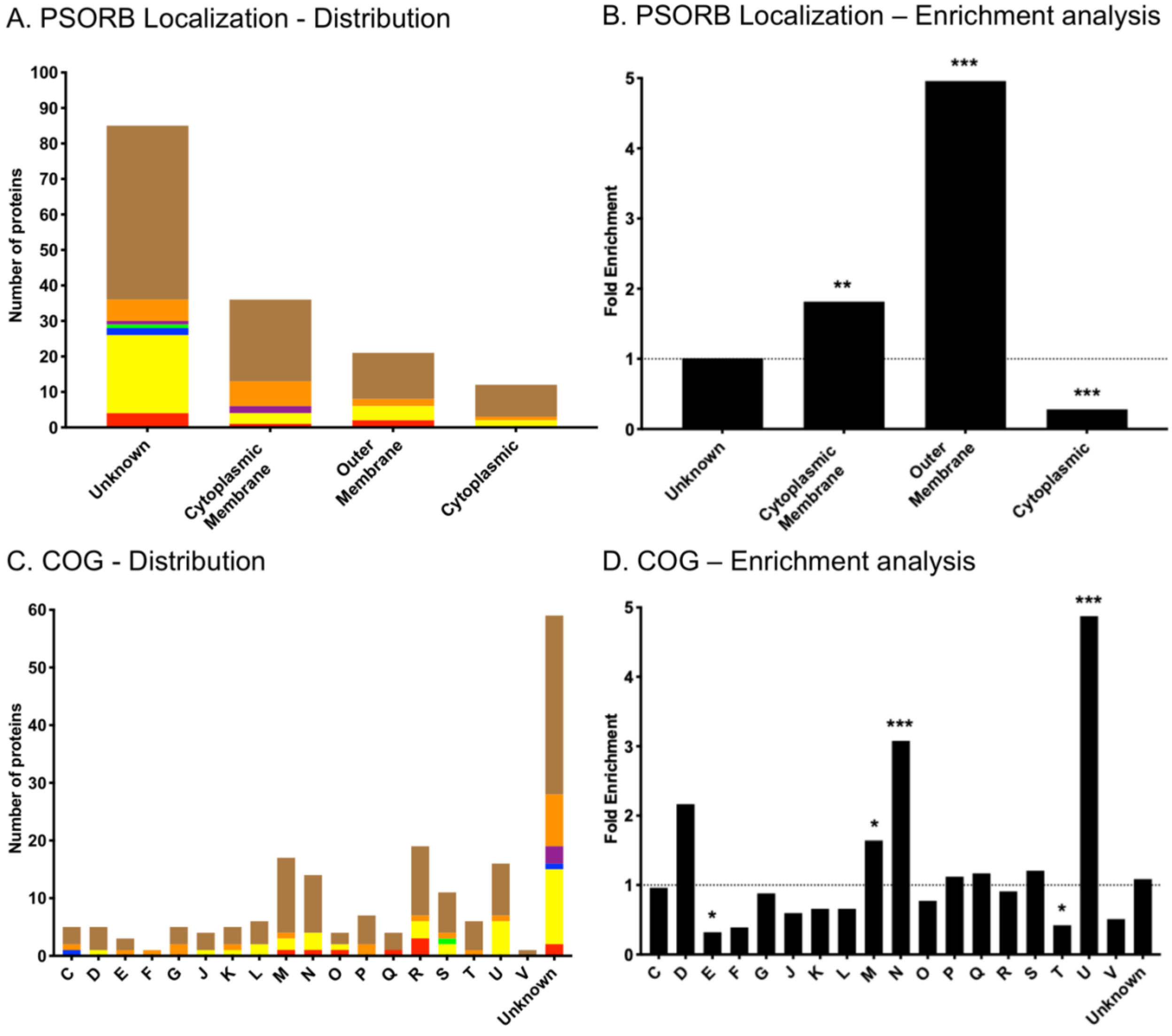

### Supplementary Figure 2

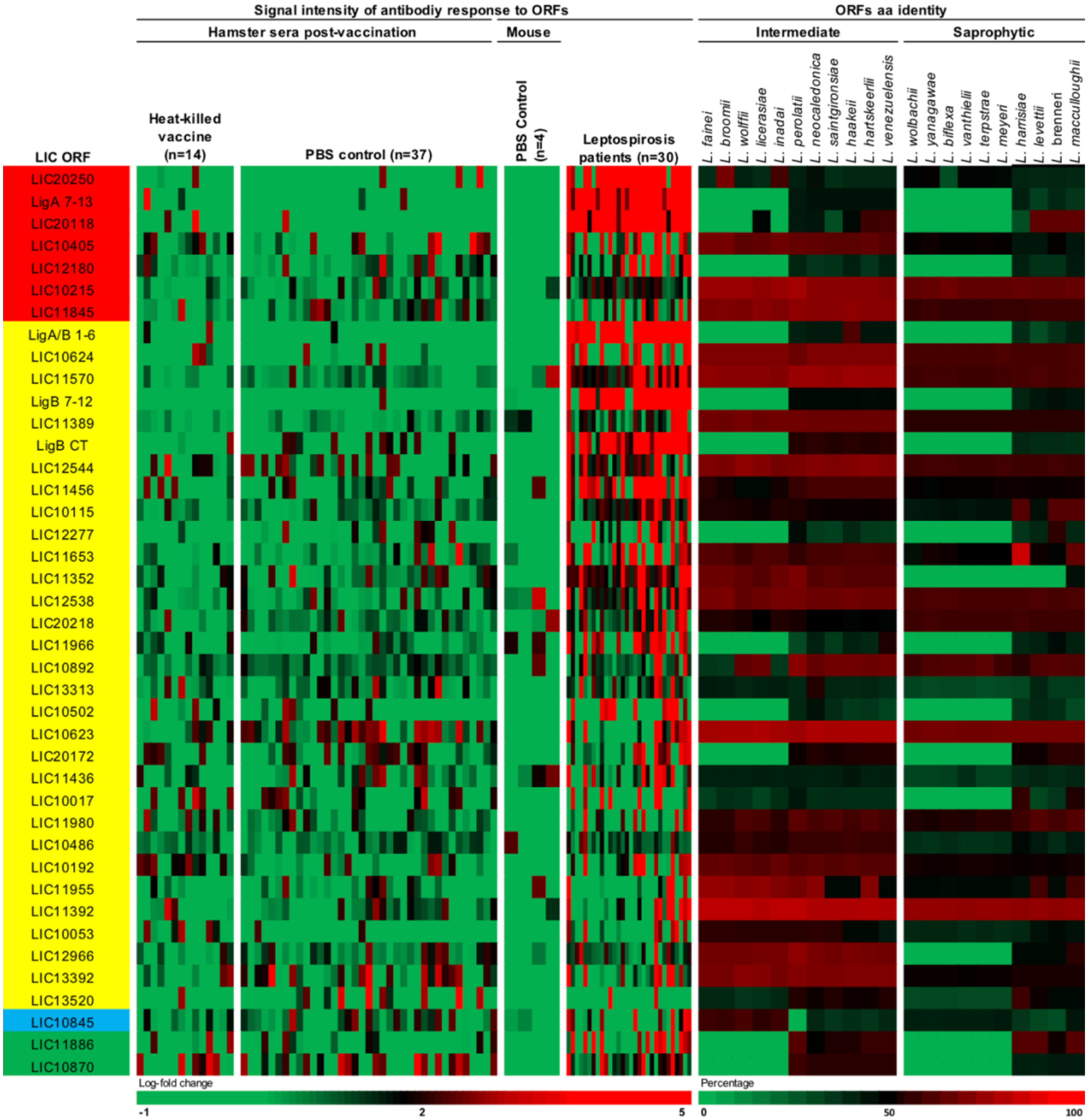

**Supplementary Figure 3**

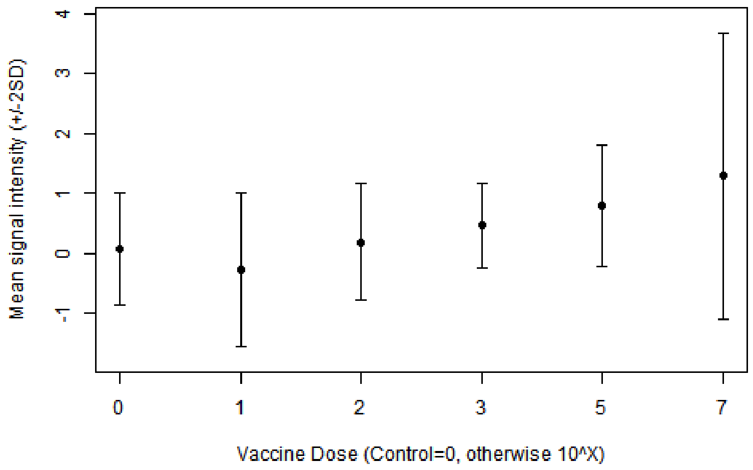

**Figure S1. Related to Figure 5. In-silico analysis of protein targets**. Using PSORB information and Genoscope database, we classified the 154 protein targets identified in this study by their putative localization in the cell (A) and their COG classification (C). The groups of proteins are classified as follow: proteins identified in all three groups analyzed (7, red); proteins identified in both hamster and mouse vaccinated with a dose of $10^7$ leptospires of the attenuated-vaccine (31, yellow); proteins identified between the group of mice immunized with different doses and the group of mice immunized with a dose of $10^7$ leptospires (2, green); proteins identified between the group of mice immunized with different doses and the group of hamsters immunized with a dose of $10^7$ leptospires (1, blue); proteins identified only in the group of mice immunized with different doses (3, purple); proteins identified only in the group of mice immunized with a dose of $10^7$ leptospires (16, orange); and proteins identified only in the group of hamsters immunized with a dose of $10^7$ leptospires (94, brown). We also performed enrichment analysis of the reactive antigens compared to the whole proteome of *Leptospira*, based on the PSORB localization (B) and COG (D). Statistical results are represented by * ($p<0.05$), ** ($p<0.001$) and *** ($p<0.0001$). See Table S4.

**Figure S2. Related to Figure 5. Complementary heat-map of 41 seroreactive proteins recognized by hamsters and mice immunized with attenuated L1-130 FcpA$^-$ attenuated-vaccine.** Proteins were selected based on the groups depicted on Figure 4 and Table S4: present in all three groups of analysis (red), present in both hamster and mice immunized with $10^7$ leptospires (yellow), present in both hamsters immunized with $10^7$ leptospires and mice immunized with a dose range (blue), and present in both mice immunized with $10^7$ leptospires and mice immunized with a dose range (green). The proteins are identified by their *L. interrogans* serovar Copenhageni ORF number and the heat-map shows the signal intensity of antibody response (based on log-fold change) in all control animals used for this analysis (14 hamsters vaccinated with heat-killed vaccine, 37 PBS control hamsters and 4 PBS control mice). The heat-map also shows the result for 30 leptospirosis patients. Right panel shows amino acid sequence identity of respective ORFs among a representative of all intermediate and saprophytic *Leptospira* species. See Figure 5.

**Figure S3. Related to Figure 4. Mouse Dose Response Relationship.** Analysis showing an association between the different doses of the attenuated L1-130 FcpA$^-$ attenuated-vaccine in mice and the mean signal response intensity against all different proteins.
