## Supplementary tables for "A Live Attenuated Vaccine Model Confers Cross-Protective Immunity Against Different Species of *Leptospira* spp."

**Supplementary Table 1.** Leptospira strains used for challenge after vaccination with L1-130 *fcpA*- mutant

| Species | Serovar | Strain | $LD_{50}$ | |
|---|---|---|---|---|
| | | | **Intraperitoneal** | **Conjunctival** |
| *L. interrogans* | Copenhageni | Fiocruz L1-130 | <10 | $2.15 \times 10^6$ |
| | Canicola | Kito | <10 | $1.78 \times 10^7$ |
| | Pomona | PO-06-047 | <10 | $<10^8$ |
| | Manilae | L495 | <10 | $2.15 \times 10^7$ |
| *L. kirschneri* | Grippotyphosa | RM-52 | $<10^3$ | $5 \times 10^7$ |
| *L. borgpetersenii* | Hardjo-bovis | 203* | ND | $>10^8$ |
| | Hardjo-bovis | J197 | $10^4$ | $<10^8$ |

* This strain was described to cause only kidney colonization in hamsters following intraperitoneal (IP) infection [17], for that reason there was no $LD_{50}$ for IP route. However, in two experiments the fatality rate was 25% after infection with $10^8$ leptospires by conjunctival (CJ) infection.

**Supplementary Table 2.** Efficacy of the immunization with attenuated *fcpA*$^-$ mutant in hamsters after challenge with $10^8$ leptospires with homologous or heterologous strains by conjunctival route

| | Challenge | | | No. | No. | Vaccine Efficacy (%, 95%CI)[¶] | |
|---|---|---|---|---|---|---|---|
| **Vaccine[£]** | **Species** | **Serovar** | **Strain** | **Expt.** | **Animals** | **Death** | **Colonization** |
| *fcpA*$^-$ | *L. interrogans* | Copenhageni | L1-130 | 4 | 6, 7, 9, 9 | 100 (90.4–100) | 80.6 (63.4-91.2) |
| | | Manilae | L495 | 3 | 7, 9, 9 | 100 (88.4–100) | 20 (8.4-39.6) |
| | | Pomona | Pomona | 2 | 8, 8 | 100 (82.9-100) | 0 (0–17.1) |
| | | Canicola | Kito | 2 | 7, 8 | 100 (82-100) | 26.7 (10.5-52.4) |
| | *L. kirschneri* | Grippotyphosa | RM52 | 2 | 7, 7 | 100 (80.9-100) | 34 (19.6-58.7) |
| | *L. borgpetersenii* | Hardjo-bovis | JB197 | 2 | 7, 7 | 100 (80.9 – 100) | 0 (0-19.8) |
| | | Hardjo-bovis | 203 | 2 | 7, 7 | 100 (47-100)[§] | 35.7 (16-61.4) |
| Heat-killed | *L. interrogans* | Copenhageni | L1-130 | 2 | 8, 9 | 58.9 (36-78.4) | 35.3 (17.2-56.4) |
| | | Manilae | L495 | 2 | 8, 9 | 5.6 (0-27.7) | 0 (0–15.5) |

[£] *Leptospira interrogans* serovar Copenhageni strain L1-130
[¶] Calculations based on frequency of outcomes compared to PBS-immunized animals
[§] Only 3/14 animals were euthanized due to clinical signs within the control group

**Supplementary Table 3.** Efficacy of the immunization with attenuated *fcpA*$^-$ mutant in mice after challenge with $10^8$ leptospires of heterologous strain by intraperitoneal route

| **Vaccine**[£] | **Vaccine Dose**[*] | **No. Experiment** | **No. Animals** | **Vaccine Efficacy (%, 95%CI)**[¶] | |
|---|---|---|---|---|---|
| | | | | **Death** | **Colonization** |
| *fcpA*$^-$ | $10^7$ | 3 | 8, 6, 4 | 100 (84.5–100) | 100 (84.5-100) |
| | $10^5$ | 2 | 4, 4 | 100 (70.7–100) | 50 (21.5-78.5) |
| | $10^3$ | 2 | 4, 4 | 100 (70.7-100) | 25 (6.3–59.9) |
| | $10^2$ | 2 | 4, 4 | 50 (21.5-78.5) | 0 (0-29.3) |
| | $10^1$ | 2 | 4, 4 | 0 (0-29.3) | 0 (0-29.3) |

[*] *Leptospira interrogans* serovar Manilae strain L495
[£] *Leptospira interrogans* serovar Copenhageni strain L1-130
[¶] Calculations based on frequency of outcomes compared to PBS-immunized animals
